# Rapid statistical methods for inferring intra- and inter-hospital transmission of nosocomial pathogens from whole genome sequence data

**DOI:** 10.1101/442319

**Authors:** Marianne Aspbury, James Sciberras, Jukka Corander, Sion C. Bayliss, Tjibbe Donker, Edward J. Feil, Richard James

## Abstract

Whole genome sequence (WGS) data for bacterial pathogens can provide evidence as to the source of nosocomial infection, and more specifically the ability to distinguish between intra- and inter-hospital transmission. This is currently achieved either through using SNP thresholds, which can lack statistical robustness, or by constructing phylogenetic trees, which can be computationally expensive and difficult to interpret. Here we compare two alternative statistical approaches using 1022 genomes of methicillin resistant *Staphylococcus aureus* (MRSA) clone ST22. In 71% of cases both methods predict the same hospital origin, which is also supported by the ML tree. Robust assignments are divided approximately equally between intra-hospital transmission and inter-hospital transmission. Our approaches are rapid and produce intuitive output that could inform on immediate infection control priorities, as well as providing long-term data on inter-hospital transmission networks. We discuss the strengths and weakness of our methods, and the generalisability of this approach.

**One Sentence Summary:** We present rapid statistical methods for distinguishing intra- versus inter-hospital transmission of bacterial pathogens using whole genome sequence data; these methods do not require the use of SNP thresholds or the generation and interpretation of phylogenetic trees.

## Introduction

Whole-genome sequencing (WGS) has proven utility for molecular epidemiological surveillance of nosocomial pathogens, and provides the power to track the emergence and spread of high-risk clones within single health-care settings, or on regional, national or global scales *(1–7)*. The recent development of community-oriented database platforms, analytical methods and data visualisation tools *(8, 9)* provide the critical infrastructure for WGS data to assume a pivotal role in guiding infectious disease management policies. These platforms provide powerful community resources for epidemiological surveillance and detailed retrospective studies. However, significant challenges remain with respect to reconstructing local transmission pathways and source attribution within a time frame that can inform on immediate infection control priorities. There is an urgent need for the establishment of a national platform based on WGS data that can provide real-time statistical probabilities of intra-hospital transmission indicative of an emerging outbreak, versus inter-hospital transmission via staff movement or patient referral *(10, 11)*.

Conventional epidemiological information such as date of infection, a patient’s current or past ward / hospital, and detailed clinical data continue to underpin infection control efforts. However, the use of large representative reference WGS databases, combined with local capacity for routine high throughput sequencing, provides substantial additional power to resolve transmission networks and define outbreaks *(5, 12)*. Currently, epidemiological inference from WGS data almost always involves the reconstruction of a phylogenetic tree; for recent examples see *(5, 13)*. Whilst this is an established approach for retrospective studies, the utility of building and interpreting trees for real-time infection control is more limited. In addition to being computationally expensive and thus potentially time-consuming, phylogenetic trees are not always statistically robust, and their interpretation is not always simple. Phylogenetic clusters are often defined on pragmatic grounds, and any given query isolate may fall awkwardly between two clusters, or else within a single cluster containing isolates from multiple other hospitals. The use of SNP thresholds for distinguishing between intra-versus inter-hospital transmission offers a practical short-cut, but one which either provides no phylogenetic context, little statistical rigour or measures of uncertainty, or else will require regular re-evaluation in the light of rapidly expanding databases. Moreover, in those cases where SNP thresholds rule out intra-hospital transmission this approach provides no direct evidence as to the likely source of the isolate.

Here we address this critical methodological gap by proposing and validating two rapid independent statistical approaches, one heuristic and one Bayesian, each of which generate intuitive graphical output, in a standardized format, regarding the probability of intra-versus inter-hospital transmission. The methods both predict the most likely hospital origin of a query isolate by comparison to a reference database, and require no capacity or experience in building or interpreting phylogenetic trees. We validated the methods using a large (>1000 genomes) WGS dataset of the important *Staphylococcus aureus* hospital-acquired (HA)-MRSA clone ST22 (EMRSA-15), representing multiple hospitals in the UK and Ireland and isolated over a ten year window (2001-2010) *(2, 6, 14)*. We consider the robustness of these methods with respect to sampling evenness, intra-hospital diversity and temporal breadth.

The methods are broadly consistent with each other and with inferences based on a maximum-likelihood (ML) tree. Discrepancies between the methods either represent subtle yet important strengths and weaknesses of each method, or else a paucity of signal in the data. The analysis provides robust hospital assignments, defined as being consistent for both methods and with the ML tree, for over two-thirds of the 91 isolates tested against the reference database.

Approximately half of these robust assignments point to intra-hospital transmission, and half to inter-hospital transmission. Surprisingly, almost 40% of inter-hospital events are long-range transmissions between “referral regions”. Finally, we discuss the utility of these methods for infection control and, more broadly, in reconstructing transmission networks from large WGS datasets. We also consider the potential challenges in deploying our approach to other species and datasets.

## Results

### Assignment of intra-versus inter hospital transmission

Preliminary analyses confirmed substantial geographical structure data (see Methods). The ML tree for all 1022 isolates is given in Figure 1. The colours of the inner ring and tree branches correspond to referral clusters and the colours of the outer ring refer to hospitals. These colour assignments are consistent with all subsequent figures, and supplemental figures showing individual clades are all taken from this tree. We took each of the 91 isolates from 2010 in chronological order (earliest first) as query isolates, and used our methods to compare these against the database of all remaining isolates from 2001-2009. Once a query isolate had been analysed, it was then added to the reference database. We compared the outputs of the heuristic and Bayesian methods (see Methods) against each other, against the ML tree and against the known origin of each isolate at the level of hospital and referral cluster. We provide examples of the outputs in the supplementary material to illustrate each of the cases we discuss below (Figures S1–S8). Figure 2 summarises these comparisons. At the hospital level, 32 of the 91 query isolates (35%) gave consistent predictions for both methods, which also concurred with the position of the isolates on the ML tree, and with the actual hospital origin. Thus, about one third of the 91 isolates from 2010 are unambiguously associated with one specific hospital. At the level of referral cluster, this figure increases to almost two-thirds (n=57; 62%). In many cases the signal from the tree is unambiguous, in that the query strain is embedded within a clade consisting of isolates representing a single hospital (see example in Figure S1). However, both methods are also robust in cases where the query isolate happens to be closely related to isolates that do not correspond to the majority hospital of the other isolates in the clade, (e.g. Figure S2), or when the query is closely related to only a single isolate corresponding to the originating hospital (e.g. Figure S3).

**Figure 1:**
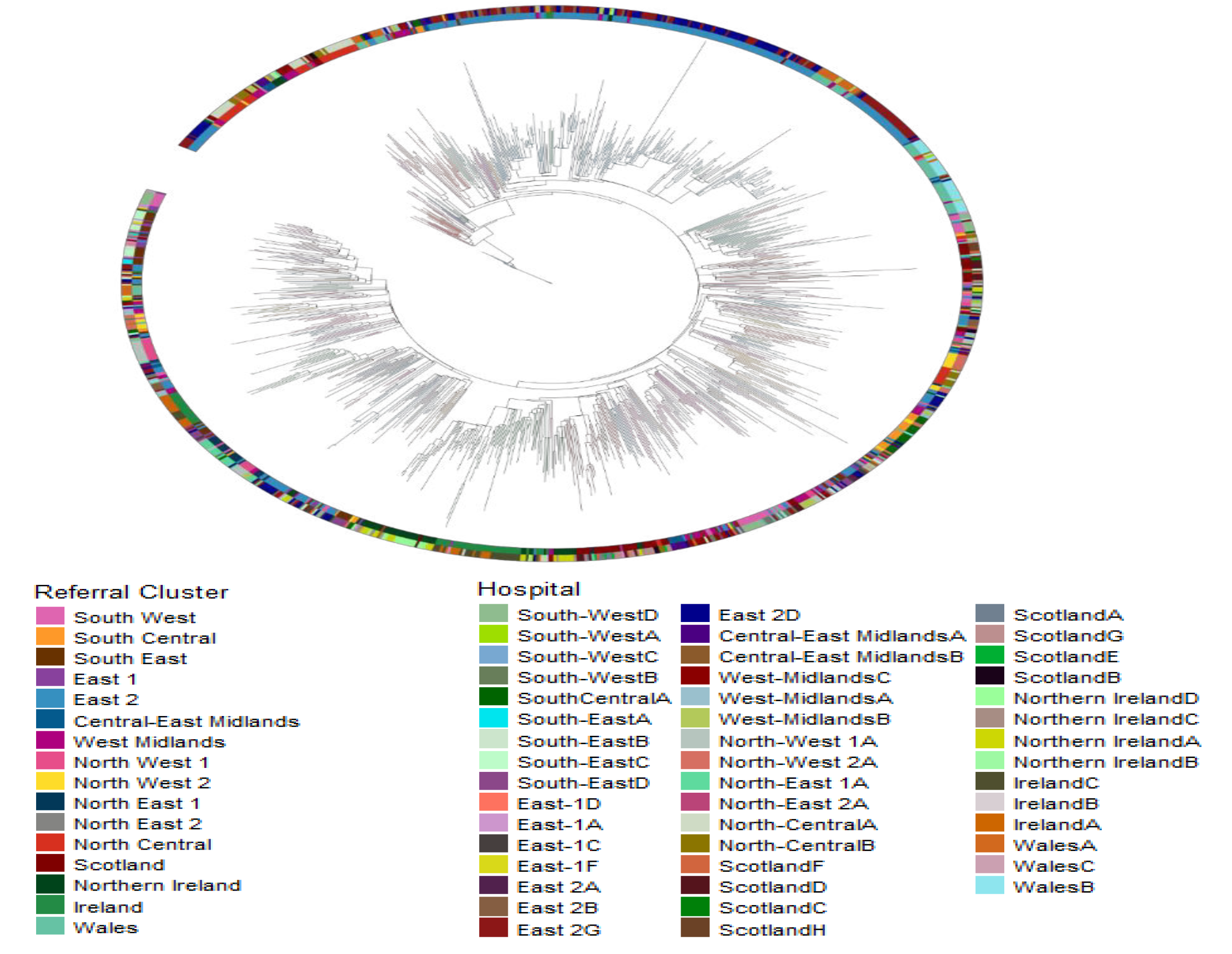
ML tree of all 1022 isolates. The colours of the inner ring and tree branches correspond to referral cluster. The colour of the outer ring corresponds to hospitals. For the positions of the referral clusters in England see Figure 4.

There are 49 cases (54%) where neither method produces an output consistent with the actual hospital origin (Figure 2). Of these, there are 33 cases (36% of total) where the two methods consistently predict the same hospital origin, but the predicted hospital is different from the actual hospital origin. These are likely to represent genuine inter-hospital transmission events. Thirteen of these 33 cases (39%) represent transmission events between referral clusters, whilst the remaining 20 cases (61%) represent transmission between hospitals in the same referral cluster. Transmission events supported by consistency between the two methods are also consistent with the phylogenetic analysis. An example of an inter-hospital transmission across referral clusters is given in Figure S4. An example of an inter-hospital transmission within a referral cluster is given in Figure S5.

**Figure 2.**
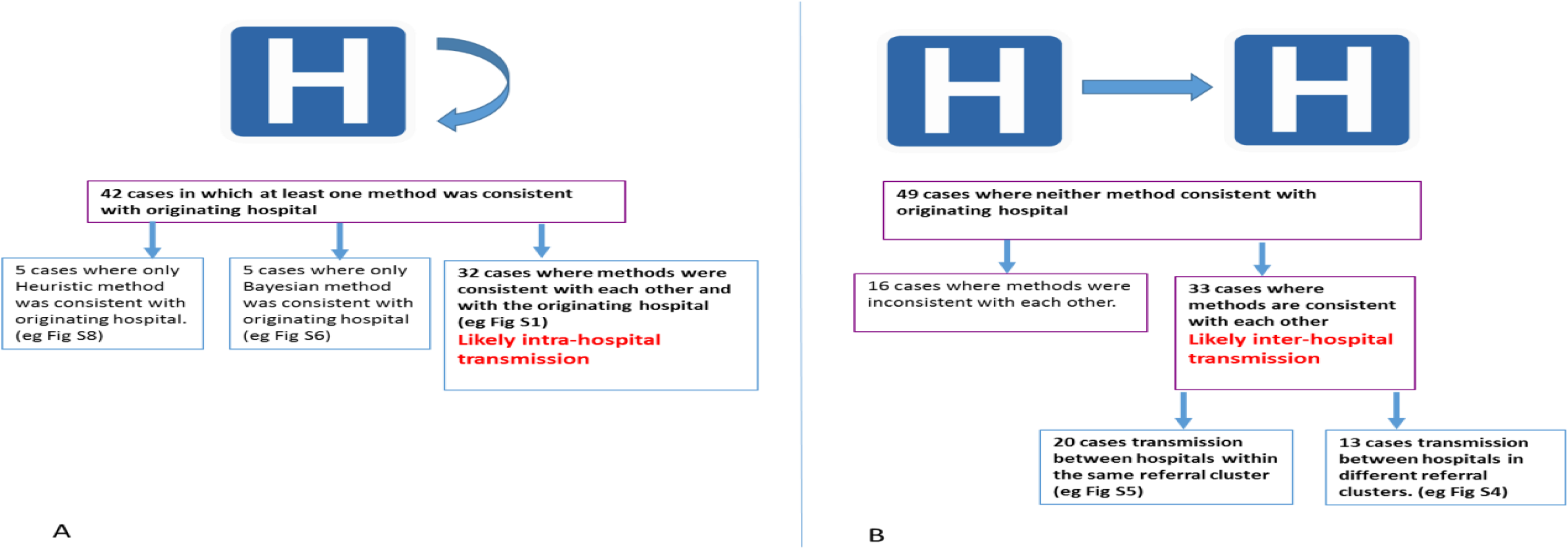
**Summary of results:** The consistency between the two methods, and the assignments where possible of intra-versus (A) versus inter- (B) hospital transmission for all 91 cases is summarised.

Thus, at the hospital level, the 91 query isolates are divided into three roughly equal groups: 35% are unambiguously associated with the hospital from which they were sampled and represent intra-hospital transmission, 36% appear to be likely inter-hospital transmission events (of which 39% represent transmission between referral clusters and 61% transmission between hospitals within the same referral clusters), and 29% are of more uncertain origin as the two methods produce inconsistent outputs. At the level of referral cluster, 62% of the isolates are clearly associated with the referral cluster from which they were sampled, 13 (14%) are likely transmission events between referral clusters (both methods agree but differ from the actual origin), and 24% are of more uncertain origin.

### Considering cases where the methods are inconsistent

In order to compare the performance of the two methods in differing scenarios, we then considered those cases where they produced inconsistent outputs. At the hospital level, there are five cases where the Bayesian output is consistent with the actual hospital origin but the heuristic output is not, and another 5 cases where the reverse is true (heuristic output is consistent with actual hospital origin but Bayesian output is not). This points to similar performance for the two methods, but indicates that each method may be more reliable under specific circumstances. At the level of referral cluster there are 10 cases where the heuristic output is consistent with the actual origin (but different from the Bayesian output) and only 3 cases where the reverse is true. In order to understand why the methods occasionally gave inconsistent results, we examined each of these cases in turn in the context of the phylogenetic tree. First, we considered the 5 cases where the output from the Bayesian method was consistent with the actual hospital origin but the heuristic method was not. In 3 of these cases, the output from the heuristic was also inconsistent with the known origin at the referral cluster level. These cases share two properties. First, the signal from the phylogenetic tree is ambiguous; that is, no single hospital is associated with isolates that are obviously closely related to the query. Second, there are very few isolates from the hospital from which the query was recovered in the reference database, but those that are present are closely related to the query. In such circumstances, the Bayesian method will predict the originating hospital on the grounds that it is more poorly represented than the alternatives, whilst the heuristic method, similar to a casual inspection of the tree, will not explicitly consider differences in sampling depth between the hospitals. An example is given in Figure S6. In this case the Bayesian method predicts the originating hospital as East1A, which is consistent with the actual origin of the query, whereas the heuristic predicts a hospital from a completely different referral cluster (SouthEastC).

Some of the isolates from 2010 represent hospitals that are not represented in earlier isolates. This means that as the query isolates from 2010 are added sequentially to the database in chronological order, there are a number of cases where the actual originating hospital of the query is not represented in the database. In these cases the heuristic cannot assign a probability to this hospital, whereas the prior implemented in the Bayesian method enables it to choose this hospital (see Methods). Thus, this represents another scenario whereby the Bayesian is more likely to predict the actual originating hospital than the heuristic method (see Fig S7 for an example).

The incorporation of sampling depth and evenness in the Bayesian method may provide some additional power in the examples above, but in other cases might produce misleading results.

This is illustrated by the 5 cases where the heuristic output is consistent with the originating hospital, but the Bayesian output is not. This can happen when a single isolate from a different hospital is somewhat related to the query isolate, a situation that might arise through a transmission event independent of the query isolate. If the hospital represented by this isolate is poorly sampled in the database, then the Bayesian method may favour this hospital even if there are isolates from more well-sampled hospitals that are more closely related to the query. If multiple distinct lineages are circulating within a well-sampled hospital, then this hospital will be even more disfavoured by the Bayesian method, as this will lower the aggregate relatedness of isolates from that hospital to the query. In contrast, the heuristic method is blind to intra-hospital diversity and will tend to simply assign the hospital corresponding to the closest isolates, regardless of the presence of more diverged sub-groups within that hospital (Figure S8).

We then considered the 8 cases where the Bayesian and heuristic methods were inconsistent with each other and also with the actual origin of the isolate at the RC level. In 4 of these cases, the position of the query isolate on the tree is consistent with an inter-hospital transmission event. Whilst both methods detect that the query isolates do not clearly correspond to the hospital (or referral cluster) from which they were sampled, they differ in their predictions of where the query isolate originated from. These differences again reflect the level of intra-hospital diversity and the difference in sampling depth between hospitals. The different scenarios are illustrated by the hypothetical trees in Figure 3. In sum, the Bayesian method will favour hospitals that are poorly represented in the database, but are represented by isolates closely related to the query isolate, over hospitals that are well sampled and contain diverse isolates, some of which are closely related to query and others of which are completely unrelated to the query (eg cases 2 and 3 in Figure 3). The heuristic will simply tend to predict hospitals represented by the isolates that are most closely related to the query, regardless of whether those hospitals also contain completely unrelated isolates or how well sampled those hospitals are.

**Figure 3.**
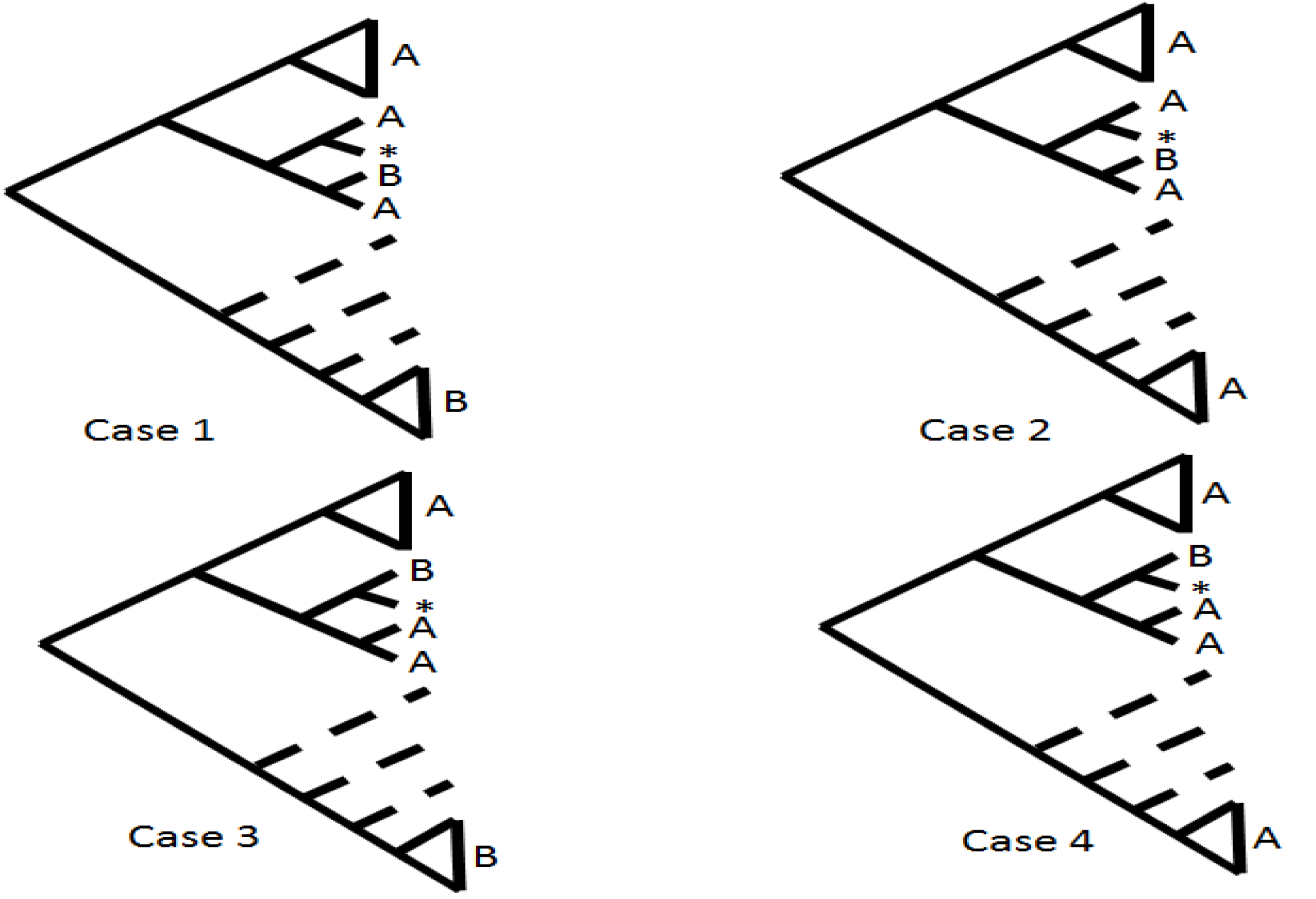
Differing scenarios that may result in inconsistencies between the methods. The position of the query is marked by an asterisk, and A and B represent two different hospitals. In case 1, both the heuristic and Bayesian methods will tend to predict hospital as the origin of the query isolate. In case 2, if hospital B is poorly sampled and unrelated genotypes are co-circulating in A, then the Bayesian method will predict B whilst the heuristic will predict A. Similarly, in case 3 the heuristic is likely to predict B, but the Bayesian method may predict A if the sample from B contains multiple genotypes. In case 4 both methods will tend to predict B.

### The temporal breadth of the database

Finally, we examined how the temporal spread of the reference database affects the predictions made by the two methods. This was motivated by the observation that those SNPs observed earlier in the sampling window also tend to be the most widely distributed among the hospitals (Figure S13). This means that older SNPs will, on average, be less informative than those SNPs that have arisen more recently. We even considered it possible that the inclusion of very old isolates in the database may result in the predictions becoming less reliable by worsening the signal to noise ratio. We therefore repeated the analysis using the 91 isolates from 2010 in turn as query isolates against the reference database, but consecutively removing the isolates from the earliest year in the database (ie first removing all the isolates from 2001, then all the isolates from 2002 and so on). In each case we recorded the fraction of the isolates in which the predictions agreed with the actual origin (at the hospital and referral cluster level), and the mean score of the predicted hospital (ie the strength of signal). Whilst this analysis does not provide any clear indication that the inclusion of older isolates significantly confounds the final predictions of the methods, the strength of signal of the predicted hospital or referral cluster is weakened with the inclusion of older isolates when using the Bayesian method (Figure S9)

## Discussion

Whole genome sequencing (WGS) has rapidly become a central tool for the molecular epidemiology of bacterial pathogens. In addition to shedding light on transmission dynamics, WGS data can provide predictions of antibiotic resistance profiles *(15, 16)* and, potentially, virulence capability *(17, 18)*, within a time-frame sufficiently short to inform on treatment options. Here we describe two statistical approaches that further exploit WGS data by distinguishing cases of infection arising through intra-hospital transmission, from those more likely to have been imported from outside the hospital (inter-hospital transmission). In 71% of the 91 cases tested the two methods were consistent, and the predicted source concurred with the phylogenetic context of the query isolate. Thus both methods perform well in cases where the phylogenetic signal is relatively strong, and the provision of intuitive output in real-time has significant implications for infection control priorities. If, in a given hospital, the majority of cases were found to be due to intra-hospital transmission, then resources should be prioritised to re-evaluating infection control procedures, and identifying putative sources of infection including individual health-care workers (HCWs). If consecutive cases of intra-hospital transmission occurred over a short period, and the isolates possessed identical “hospital profiles” from our procedures, then this would be strongly indicative of an outbreak. In contrast, if the majority of cases were imported from other hospitals, then screening on admission and/or prior to referral should be prioritised.

In addition to providing an invaluable tool for real-time epidemiology, the data produced by these methods provide useful summary statistics regarding the relative frequency of intra-versus inter-hospital transmission. Given the strong spatial structuring evident from an overall inspection of the ML tree, it might have been expected that intra-hospital transmission is much more frequent than inter-hospital transmission, but our analyses suggests this is not the case. Approximately half of the robustly assigned cases were consistent with intra-hospital transmission and half with inter-hospital transmission. Moreover, of the inter-hospital transmission events, 39% were long-range events between hospitals in different referral clusters. These observations are consistent with the recent report by Auguet et al, which also highlighted the significance of inter-hospital transmission of *S. aureus* ST22 *(19)*. Whilst all of these transmission events were corroborated by an inspection of the tree, this validation step took far longer than generating the output, and compiling summary statistics from a brute-force inspection of the tree alone would be a painstaking and subjective process. The utility of our methods to rapidly catalogue multiple transmission events could pave the way towards constructing transmission networks within and between hospitals based on the frequency of actual events. This contrasts with previous studies based on broad measures of differentiation between bacterial populations circulating in different hospitals *(14)*.

In 29% of the cases we examined, our two methods did not produce consistent results, and in these cases the phylogenetic signal tended to be more ambiguous. Although our methods appear to be reasonably robust to sampling uneveness and temporal breadth, an obvious limitation lies in the fact that the reference database does not represent all of the major hospitals in the UK and Ireland, and some hospitals are only represented by a very small number of isolates. Whereas the Bayesian method can account for cases where the originating hospital of a query isolate is not present in the reference database, the heuristic method cannot, and this can explain some of the discrepancies. An alternative possibility is that the strains did not directly derive from a hospital at all, but from the community. Coll et al have recently demonstrated frequent transmission within the community and between the community and hospitals for *S. aureus* ST22 in the East of England *(13)*. We also note that pre-existing inter-hospital transmission events present in the database might complicate the assignment of subsequent events, and that there may be rare cases that are essentially intractable regardless of the reference database used or the sophistication of the method.

Whilst presenting two alternative methods provides a useful means by which to gauge their reliability (in terms of consistency between the methods) and their relative strengths and weaknesses, ultimately the use of dual methods is sub-optimal and adds unnecessary complexity. How then to choose which method is most suitable? Whilst our results suggest comparable performance of the two methods, one possibility going forward would be to use the heuristic method to obtain the best possible priors to inform the Bayesian method. These priors could be updated at regular intervals, thus using the heuristic method to optimise the Bayesian method as the reference database grows. Larger, more representative reference databases will be generated over time, and this will mitigate many of the limitations we have identified in this study and will help to further optimise the methodology.

A major question remains as to how generalizable our approach is to other pathogenic species, or even other clones of *S. aureus*. The ability to reliably reconstruct recent transmission chains within and between hospitals is dependent upon the epidemiological characteristics of the clone/species in question. Multidrug resistant clones that transmit primarily through the healthcare system will be much more amenable to this approach than more susceptible clones that are frequently carried and transmitted asymptomatically in the community and/or commonly occupy environmental niches. Whilst transmission of MRSA ST22 has recently been shown to occur within the community in the UK, it has also been by far the most common hospital-acquired MRSA in this country over the last two decades or so *(3, 6)*, and has spread world-wide *(20)*. Other *S. aureus* clones, such as USA300, are more frequently associated with transmission within the community *(21, 22)*, and the approach as described will have more limited utility for this clone.

Other major pathogens, such as *E. coli* and *Klebsiella*, may be even less suitable due to frequent recombination, and transmission within the community and environment. However, for such species the method should still have utility when specific nosocomial antibiotic resistant clones are considered. For example, *K. pneumoniae* ST258 is a major carbapenemase producing Enterobacteriacea (CPE) clone that has clonally disseminated over a short time period throughout the Italian healthcare network *(23, 24)* and elsewhere *(25, 26)*, and preliminary analysis using an ST258 dataset confirms the usefulness of our approach for this clone (data not shown).

In sum, here we demonstrate that it is possible, and indeed desirable, to draw inferences from WGS data regarding local epidemiology without recourse to phylogenetic analysis or SNP thresholds. We believe the methods as presented provide an important first step towards a more efficient exploitation of WGS for real-time infection control, and for building a more detailed picture of the transmission dynamics of multidrug resistant clones through health-care networks.

## Materials and Methods

### The isolates

Our analysis was based on the dataset of Donker et al. *(14)* and consists of 1022 isolates of *S. aureus* ST22 recovered from cases of invasive disease from 46 hospitals from England, Wales, Scotland, Northern Ireland and the Republic of Ireland from 2001 to 2010. The majority of the isolates were collected through the British Society of Antimicrobial Chemotherapy (BSAC) bacteraemia resistance surveillance programme, supplemented with additional isolates retrospectively collected from nine hospitals in the East of England over the same time-frame *(14)*.

We grouped the English hospitals into referral clusters, as previously defined by Donker et al *(14)* on the basis of geographic proximity and patient movement, as this provides a useful additional geographical scale. For hospitals outside of England, where patient referral data is not available, we consider country or province (Scotland, Wales, Northern Ireland, Republic of Ireland) as a single referral cluster. To preserve anonymity, hospital names were coded, as described previously *(14)*. Figure 4(A) shows the borders of the English referral regions, and Figure 4(B) shows the number of isolates used in this analysis from each referral region broken down by year. There is an approximately even representation of isolates across the different years. Figure 4(C) shows the number of isolates broken down by referral region and hospital. The large excess of isolates from the East of England reflects the inclusion of isolates from the second retrospective study.

**Figure 4:**
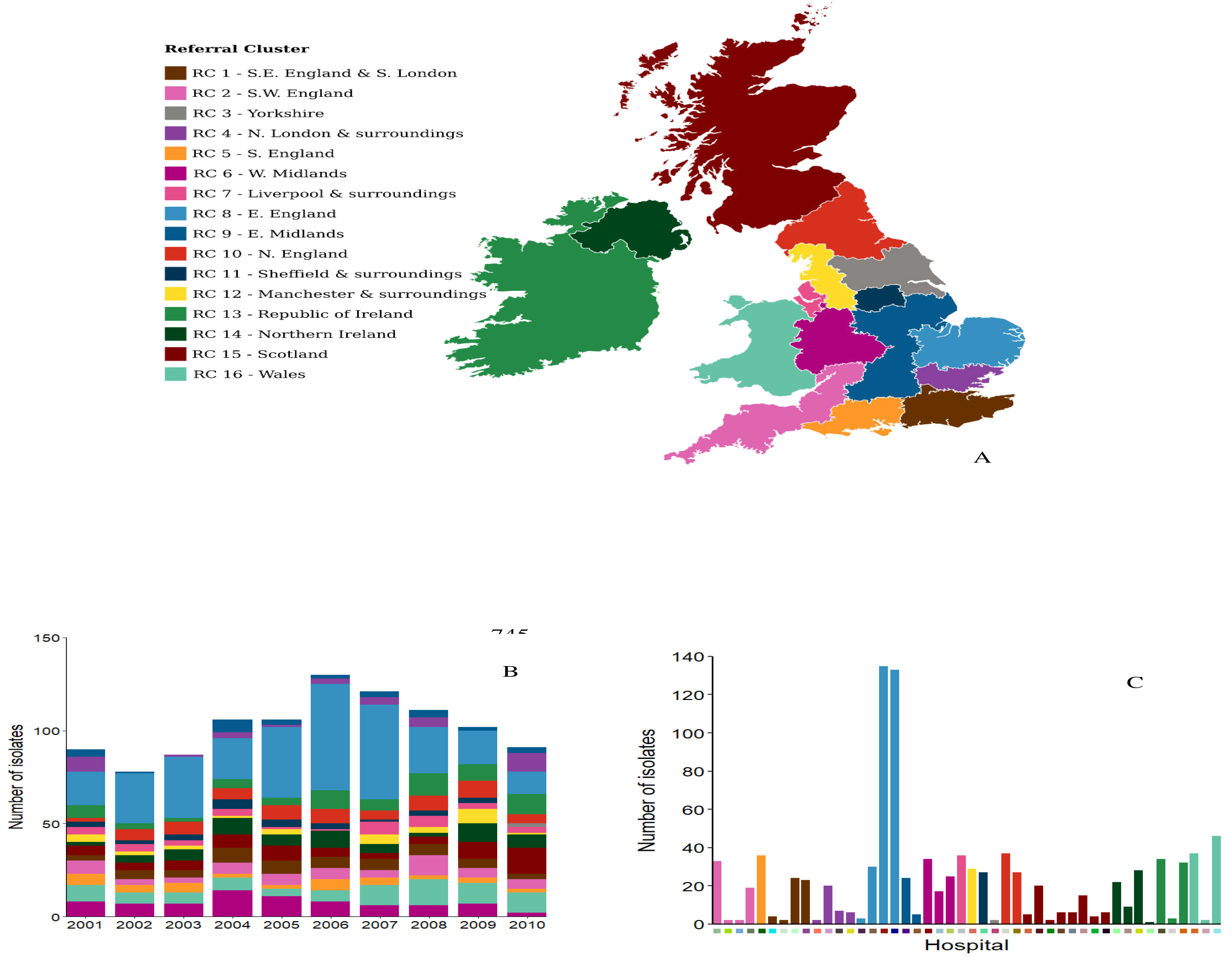
Position of the referral regions in England (A), breakdown per year in the database (B) and number of isolates in each hospital by referral region (C). The colour codes for the hospitals are given in Figure 1. Colour codes for hospitals and referral regions are used for the supplemental figures.

### The single-nucleotide polymorphisms (SNPs)

The short-read Illumina data was mapped to the ST22 reference genome MRSA252, and the regions spanning the major non-core elements (e.g. SCCmec) were excluded as described previously *(2, 14)*. We discounted all monomorphic sites, and all SNP sites where the frequency of N (ambiguous base) was >1% to ensure quality control. We also removed a small number (n =24) of SNP sites known to be associated with antibiotic resistance as these are known to be prone to homoplasy, and all tri-allelic (n = 144) and tetra-allelic (n = 4) as these may represent sequencing errors or sites under unusual selection pressure. This left a total of 23,686 bi-allelic SNPs which were carried forward to downstream analyses. Of these, 18,217 were singletons where the minority base is only present in a single isolate, and 5469 were present in >1 isolate, with the most common appearing in 454 of the 1022 isolates (Figure S10). Although singleton SNPs are not phylogenetically informative, we retained them in the reference database as it is possible they will be observed in query isolates, and thereby provide additional signal. The positions of the non-singleton SNPs are given in Figure S11; there is no indication that they are non-randomly clustered within the alignment. The mean number of SNPs per isolate was 53.7 (range 2-131).

### Preliminary analyses

We carried out some preliminary analysis to examine the degree of geographical clustering of the SNPs. A maximum likelihood tree based on the SNP data confirmed that the ST22 isolates are clustered at the levels of hospital, referral region and country (Figure 1). We further examined the degree of clustering on the hospital level by plotting, for each SNP, the number of isolates in which that SNP was found against the number of hospitals. We compared these data with a null distribution based on the number of hospitals in which each SNP is expected to occur, given their frequency, under the assumption that SNPs are randomly distributed among the hospitals (Figure S12). This analysis revealed that SNPs tend to be distributed among far fewer hospitals than would be expected by chance, thus confirming a strong geographical signal within the SNP data. However, as discussed by Donker *et al*, numerous inter-hospital transmission events are still evident from the phylogeny *(14)*.

Previous work has shown that the HA-MRSA ST22 (EMRSA-15) clone emerged from a point source in the English west-midlands in the mid-1980s *(3)*. Continued expansion, combined with occasional long-distance transmission, throughout our sampling period (2001-2010) explains why older SNPs are more common, thus likely to be more widely distributed throughout the health-care network (Figure S13). This implies that the time point at which SNPs are first observed should inversely correlate with the strength of the discriminatory signal for predicting hospital origin; that is, more recent SNPs will be restricted to fewer hospitals and will therefore be more informative.

### The statistical methods

Having carried out a preliminary analysis of a quality controlled dataset, we then developed and tested statistical methodologies for predicting hospital origin. For the English hospitals we also considered referral cluster, and for isolates from Scotland, Wales, The Republic of Ireland and Northern Ireland we considered these national regions as the higher order geographical unit. Our approach was to consider each of the 91 isolates from 2010 as query isolates. Each of these isolates were compared in turn chronologically against the reference database consisting of all isolates from 2001-2009 plus all previously compared 2010 isolates. The predictions were then compared to each other, against the context of the isolates in the maximum likelihood tree, and against the actual origin of the isolates. For each comparison we considered origin both at the level of hospital and referral region / country.

To describe each method we labelled hospitals with an index *h* = 1,…, *H*, referral clusters with an index *r* = 1,…, *R* and variable sites with an index *s* = 1,…, *S*. The reference database of *N* isolates was stored (Fig. S2) as an *S* × *N* binary matrix *<u>D</u>* whose element *D_si_* is 1 if there was a SNP (the rarer nucleotide) at site *s* in isolate *i* and is 0 otherwise. The same coding was used for the query isolate, stored in the binary *S* × 1vector *<u>x</u>*, containing 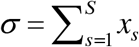 SNPs. To compare *<u>x</u>* with the *n_h_* isolates in the database that were collected from hospital *h*, we analysed the *S* × *H* matrix of SNP counts *<u>C</u>*, whose element *C_sh_* is the number of isolates from *h* that have a SNP at site *s* (0 ≤ *C_sh_* ≤ *n_h_*). In each classifier we compared *<u>x</u>* with *<u>C</u>* to assign a score *y_h_* indicating the probability that *<u>x</u>* comes from (or is consistent with isolates taken from) hospital *h*. In principle, the only job of a classifier is to find the class *ĥ* which has the largest score *y_h_*, but we will present and analyse the full set of scores to assess and compare our methods. When the classes of interest were referral clusters, not individual hospitals, we further aggregated isolates in the database to produce a *S* × *R* count matrix whose element 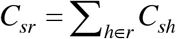, and assigned scores *y_r_* to each of the *R* referral clusters. For clarity we present detailed methods only for the case of classifying to the level of hospital.

In all our analyses the 91 query isolates from 2010 were classified in chronological order. Once classified an isolate became part of the database, by adding *<u>x</u>* as an extra column to the database *<u>D</u>*, and associating *<u>x</u>* with whichever hospital and referral cluster it was collected from. The size of the database, and the number of classes, therefore increased as we tested the 2010 isolates. At the outset, *N* = 932, *H* = 36, *R* = 15; by the end *N* = 1021, *H* = 46, *R* = 16, as in 2010 there were isolates from 10 hospitals new to the database and from one new referral cluster. The number of variable sites remained constant (*S* = 23, 686) throughout our analysis.

### Method 1: A simple heuristic framework (“heuristic”)

We first developed a simple heuristic approach by weighting the presence of a SNP in a given hospital according to the inverse of its overall frequency, reasoning that any classifier giving equal weight to all sites might be prone to having the most useful information swamped by a mass of less differentiating information.

This was achieved by building a classifier that scores only the *σ* sites that are SNPs in the query isolate *<u>x</u>* and allocates more weight to SNPs previously seen in few hospitals. We scaled the SNP counts *C_sh_* (at those sites at which *x_s_* = 1) by the total number of isolates in the database that have a SNP at site *s* to produce a measure, 
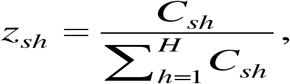
of the localisation of a given SNP. A SNP that has only ever appeared in one hospital scores a maximum *z_sh_* = 1 for that hospital, and 0 for all other hospitals. A SNP seen in equal numbers of isolates in two hospitals would produce a *z*-score of 0.5 for each of those hospitals, and so on. A SNP seen in many hospitals will produce a low *z*-score for that SNP in each of those hospitals.

The total weight of *z*-scores for each SNP is 1, so the normalised hospital score for this heuristic classifier is simply

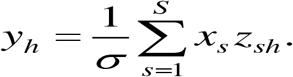

[The *x_s_* appears in this expression to ensure that only SNPs present in *<u>x</u>* are counted.]

### Method 2: A Bayesian framework (“Bayesian”)

Our second approach uses conventional naïve Bayes classification *(27)*, in which the hospital scores are posterior probabilities *y_h_* = *L_h_ P_h_* / Ω, comprising a likelihood function *L_h_* (the probability that query isolate *<u>x</u>* contains the 0s and 1s it does, given that it comes from *h*) and any prior expectation *P_h_* we have that *<u>x</u>* comes from *h*. The normalisation constant Ω = ∑*_h_L_h_P_h_*.

As there are only two possible states at each site (SNP or non-SNP) we used a Binomial likelihood function, with a Beta prior (hyper-parameters *a* and *b*) to estimate the probability that an isolate from hospital *h* has a SNP at site *s* as

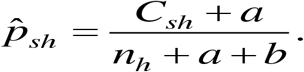

Following Gelman et al *(28)* we used the database itself to fix the hyperparameters *a* = 0.045 and *b* = 19.8 by fitting to the mean and variance of the observed distribution of the number of SNPs per site (Figure S1). The estimated probability that there is not a SNP at site *s* in hospital *h* is (1– *p*̂_*sh*_). These two probabilities are sufficient to construct the likelihood function *L_h_*. In a naïve Bayes classifier, each of the contributing pieces of evidence (sites in our case) is treated as independent, so our likelihood function is 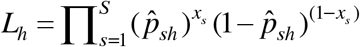.

To help us interpret the level of support for a given classification we introduced a dummy class *h* = 0 into the database. This comprised a single, invented, isolate, with a typical number of SNPs (50), none of which aligns with any of the SNPs in the database. The likelihood *L*_0_ is then easily computed, as there are 0 SNPs in *S* sites, and all of the 50 ‘extra’ SNPs are mismatches. All of the *H* true hospitals have their likelihood *L_h_* adjusted for the presence of the dummy class by the addition of 50 ‘extra’ non-SNP matches.

In the Bayesian classifier presented here *P_h_* is the product of two prior distributions. The first reflects that fact that there is uneven sampling of the hospitals in the database (Figure 3), so we might expect that a randomly-chosen query isolate is more likely to come from a highly-sampled hospital. The second prior is a simple attempt to account for the known geographical clustering of patient referrals and the MRSA genome in this dataset. Donker et al. *(14)* estimated the number of ‘candidate introductions’ - isolates that look out-of-place in the phylogenetic tree - in the same database used here. We used the frequencies of these candidate introductions to parameterise a 3-level distribution that asserts our prior expectation that the mostly likely hospital (and referral cluster) for a query isolate is the hospital (referral cluster) it was taken from. We note in passing that for most query isolates, neither of our priors makes much difference to the classification; by far the biggest source of contrast between hospitals comes from the likelihood function. The exception is the situation where the query isolate comes from a hospital that is new to the database, as happens 10 times in our test. In such cases we set the baseline score either to that of the dummy hospital, or to the likelihood *L_r_* of the origin referral cluster if there are already isolates in the database from *r*.

For practical reasons we computed the log-likelihoods 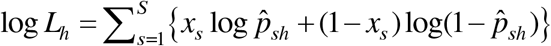 and display the Bayesian log-scores log *y_h_* = log *L_h_* + log *P_h_* – logΩ. All methods were implemented in R and output was generated for each individual query in seconds on a standard laptop.

All scripts are readily available at https://github.com/marianne-aspbury/hospital-origin.

## Acknowledgements

The ST22 data was generated as part of the UKCRC Translational Infection Research Initiative and the Medical Research Council (grant no. G1000803) with contributions to the grant from the Biotechnology and Biological Sciences Research Council (BBSRC), the National Institute for Health Research on behalf of the Department of Health, and the Chief Scientist Office of the Scottish Government Health Directorate (to Sharon Peacock). EJF is supported by the JPI-AMR project SpARK with funding contributions from the Medical Resarch Council (MR/R00241X/1), and the EPSRC (GCRF) project ReNEW (EP/P028403/1). MA received funding from the “STARS” programme, supported by the BBSRC, via the University of Bath Institute for Mathematical Innovation Summer Internship Scheme. JS was supported by a University of Bath studentship. TD is affiliated to the National Institute for Health Research Health Protection Research Unit (NIHR HPRU) in Healthcare Associated Infections and Antimicrobial Resistance at University of Oxford in partnership with Public Health England (PHE). Tjibbe Donker is based at University of Oxford. The views expressed are those of the author(s) and not necessarily those of the NHS, the NIHR, the Department of Health or Public Health England. Members of the East of England Microbiology Research Network are Benny Cherian (Basildon and Thurrock University Hospitals, Basildon, UK), Tony Elston (Colchester Hospital University, Colchester, UK), Richard Kent (Ipswich Hospital, Ipswich, UK), John Hayward (James Paget University Hospital, Great Yarmouth, UK), Louise Teare (Mid Essex Hospital Services, Chelmsford, UK), Juliet Foweraker (Papworth Hospital, Papworth Everard, UK), David Enoch (Peterborough and Stamford Hospitals, Peterborough, UK), Nada Elhag (Southend University Hospital, Westcliff-on-Sea, UK), Prema Singh (West Hertfordshire Hospitals, Watford, UK), and Rebecca Tilley (West Suffolk Hospital, Bury St Edmunds, UK).

## Author Contributions

EJF, RJ Designed the study and wrote the manuscript. MA, JS wrote the scripts, carried out the analyses and contributed to the design of the study. JC, TD contributed to the statistical methodology and SB carried out the phylogenetic analysis.

## Ethical Statement

Written informed consent from patients was not required as all bacterial isolates were collected, processed and stored as part of routine clinical care and/or bacteraemia surveillance programmes. The study protocol was approved by the National Research Ethics Service (reference 11/EE/0499), and by the Cambridge University Hospitals NHS Foundation Trust Research and Development Department (reference A092428)

## Competing Interests

We have no competing interests to declare.

We are very grateful to Nick Priest for many helpful discussion, and to Matthew Holden for excellent feedback received as part of his examination of the PhD thesis of JS. We are also grateful for helpful discussions on this work at the “Permafrost” workshop 2018. All scripts are readily available at https://github.com/marianne-aspbury/hospital-origin.

## List of supplementary material

**Supplementary Figures S1-S8** comparative examples of the outputs from both statistical methods (see text and Figure 2).

**Supplementary Figure S9** examining the impact of changing the temporal breadth of the database.

**Supplementary Figures S10-S13** preliminary analyses of the data (see Methods).

**Figure S1.**
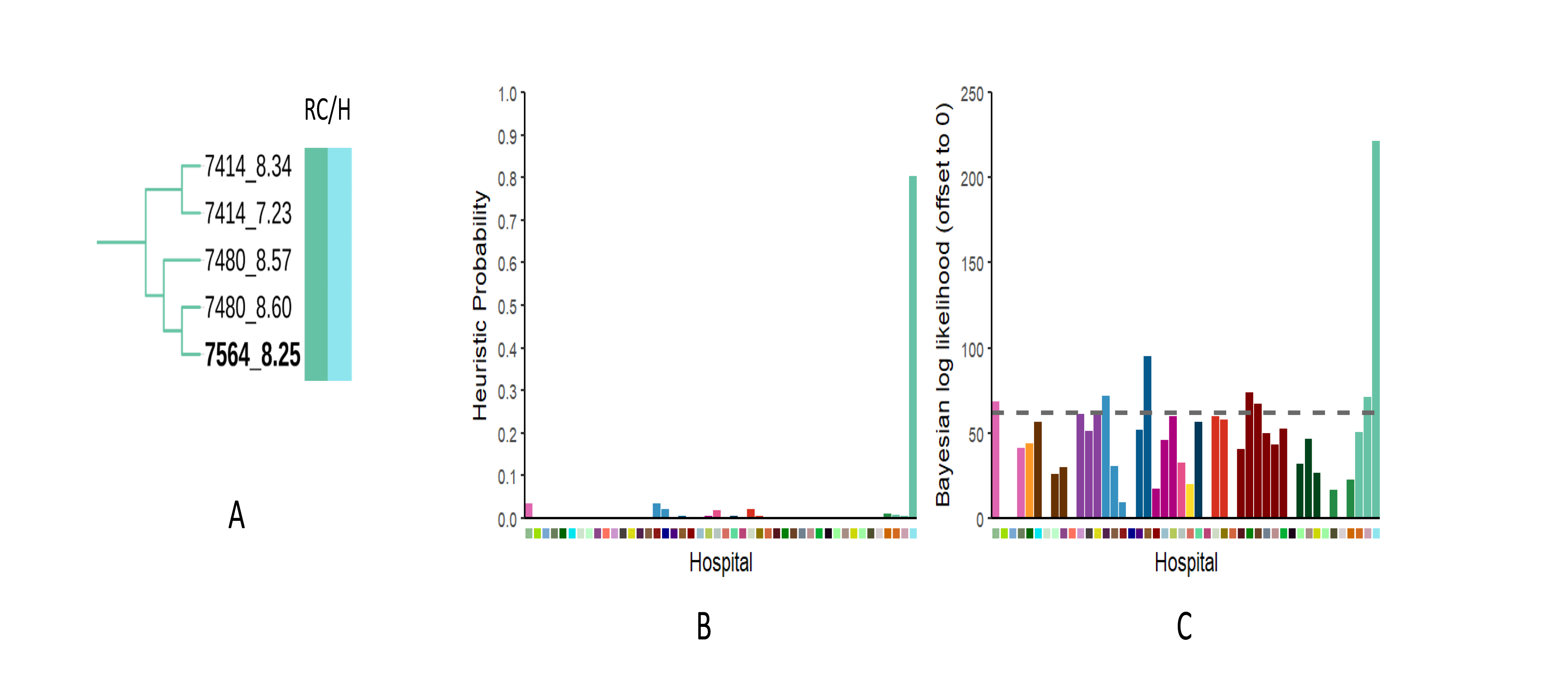
The query isolate id (in bold) is 7565_8.25 was isolated from hospital WalesB (RC Wales). Inspection of the Tree (A) confirms that it falls within a clade of other isolates from the same hospital. This hospital was also the Predicted source using both the heuristic (B) and Bayesian (C) methods. RC = Referral cluster. H = Hospital. For a key to the colours see Figure 1.

**Figure S2.**
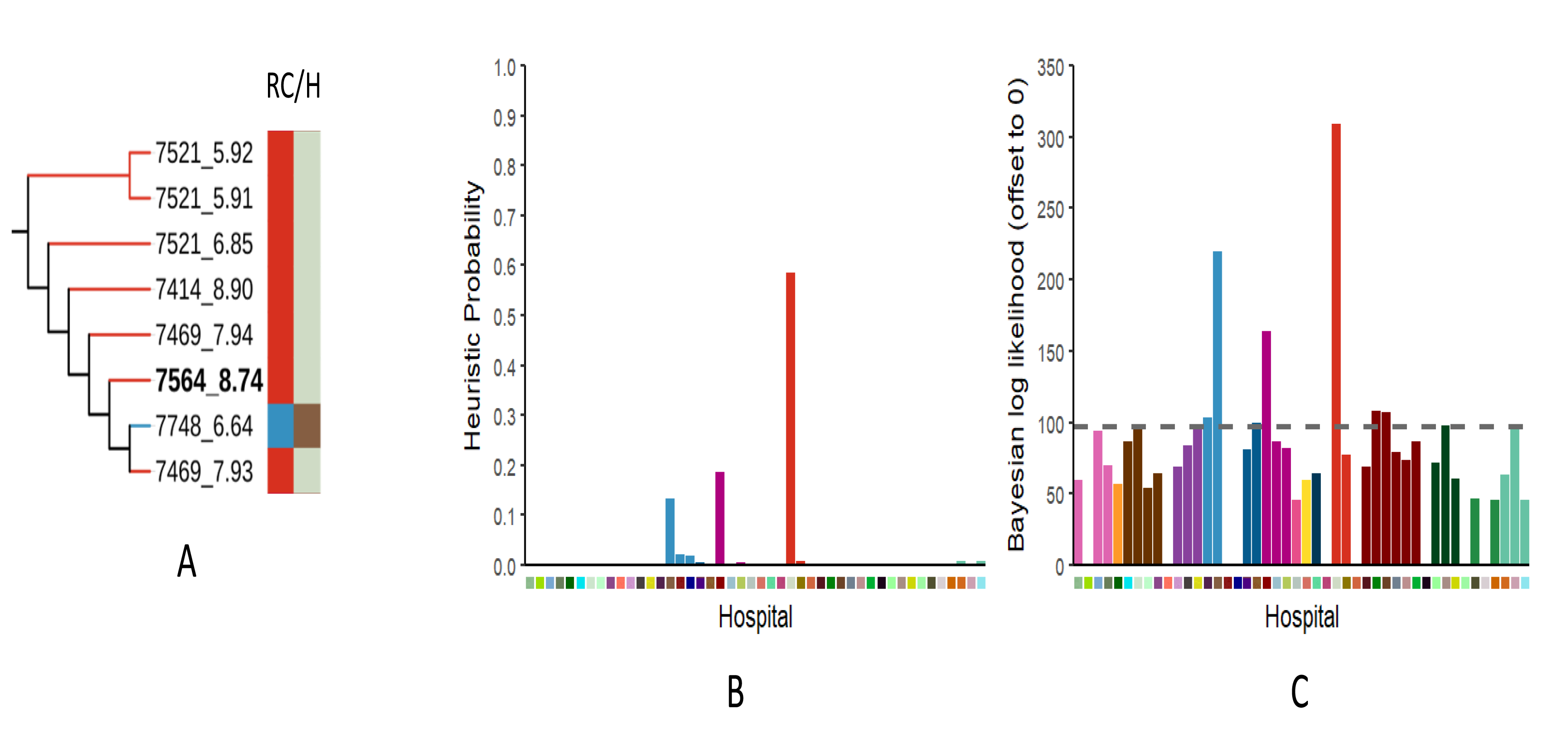
The query isolate id (in bold) is 7564_8.74 was isolated from hospital NorthcentralA. Inspection of the Tree (A) confirms that it falls within a clade of other isolates from the same hospital, but is also closely related to isolate 7748_6.64 which originated from hospital East2G (presumably due to an inter-hospital transmission). Despite this, both the heuristic (B) and Bayesian (C) methods predicted NorthcentralA as the source of the query isolate. RC = Referral cluster. H = Hospital.

**Figure S3.**
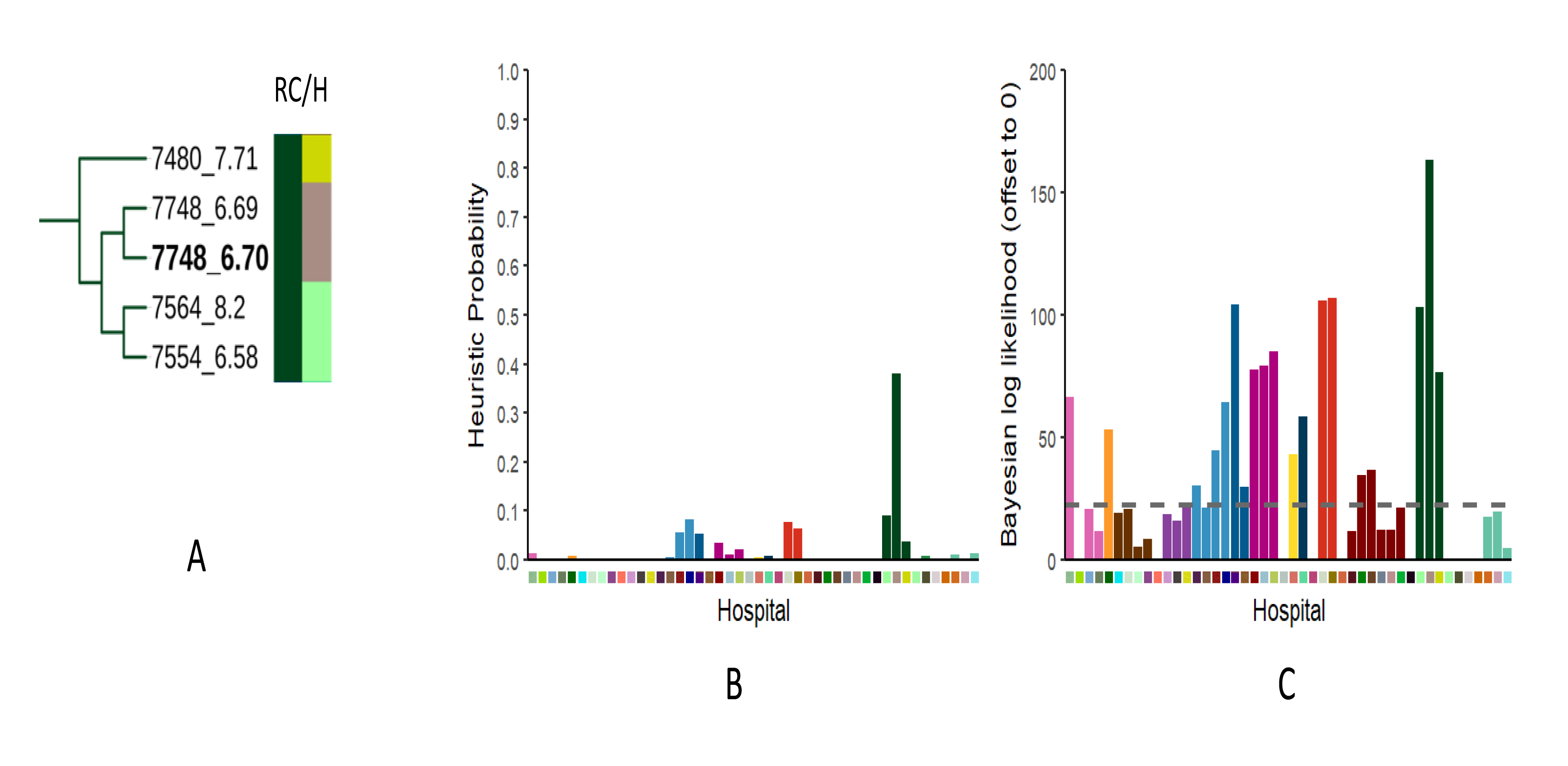
The query isolate id (in bold) is 7748_6.70 was isolated from hospital NorthernIrelandC. Inspection of the Tree (A) confirms that it is most closely related to isolate 7748_6.69 which was also isolated from hospital NorthernIreland C, but is not embedded in a large clade of isolates from this hospital. Both the heuristic (B) and Bayesian (C) methods predicted NorthernIrelandC as the source of the query isolate.

**Figure S4.**
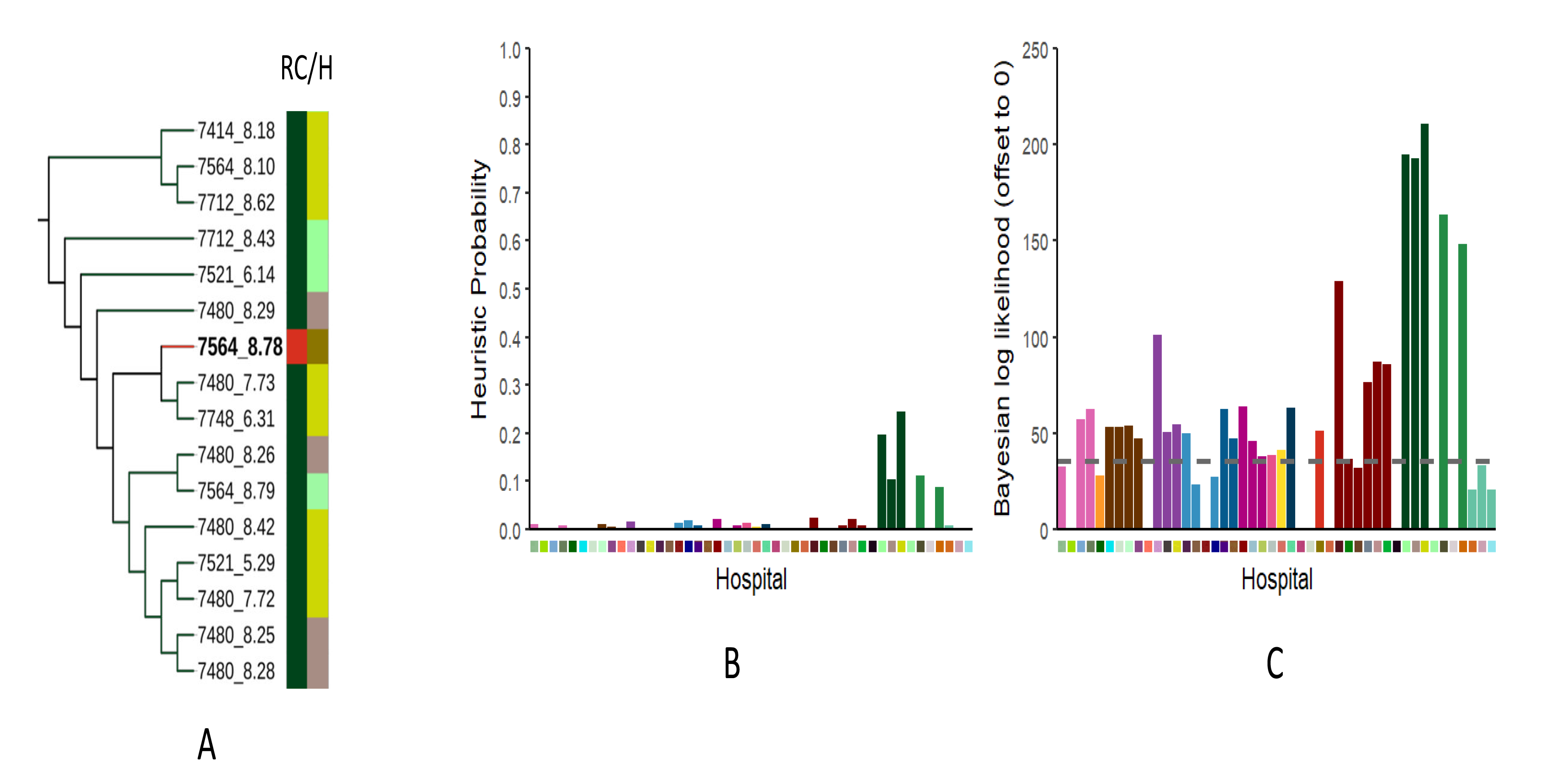
The query isolate id (in bold) is 7748_6.70 was isolated from hospital NorthCentralB. However, inspection of the Tree (A) confirms that it corresponds to a clade of isolates from Northern Ireland, indicating an inter-hospital transmission Event between referral clusters. Both the heuristic (B) and Bayesian (C) methods predicted NorthernIrelandA as the source of the query isolate, which is Consistent with inter-hospital transmission. RC = Referral cluster. H = Hospital. For a key to the colours see Figure 1.

**Figure S5.**
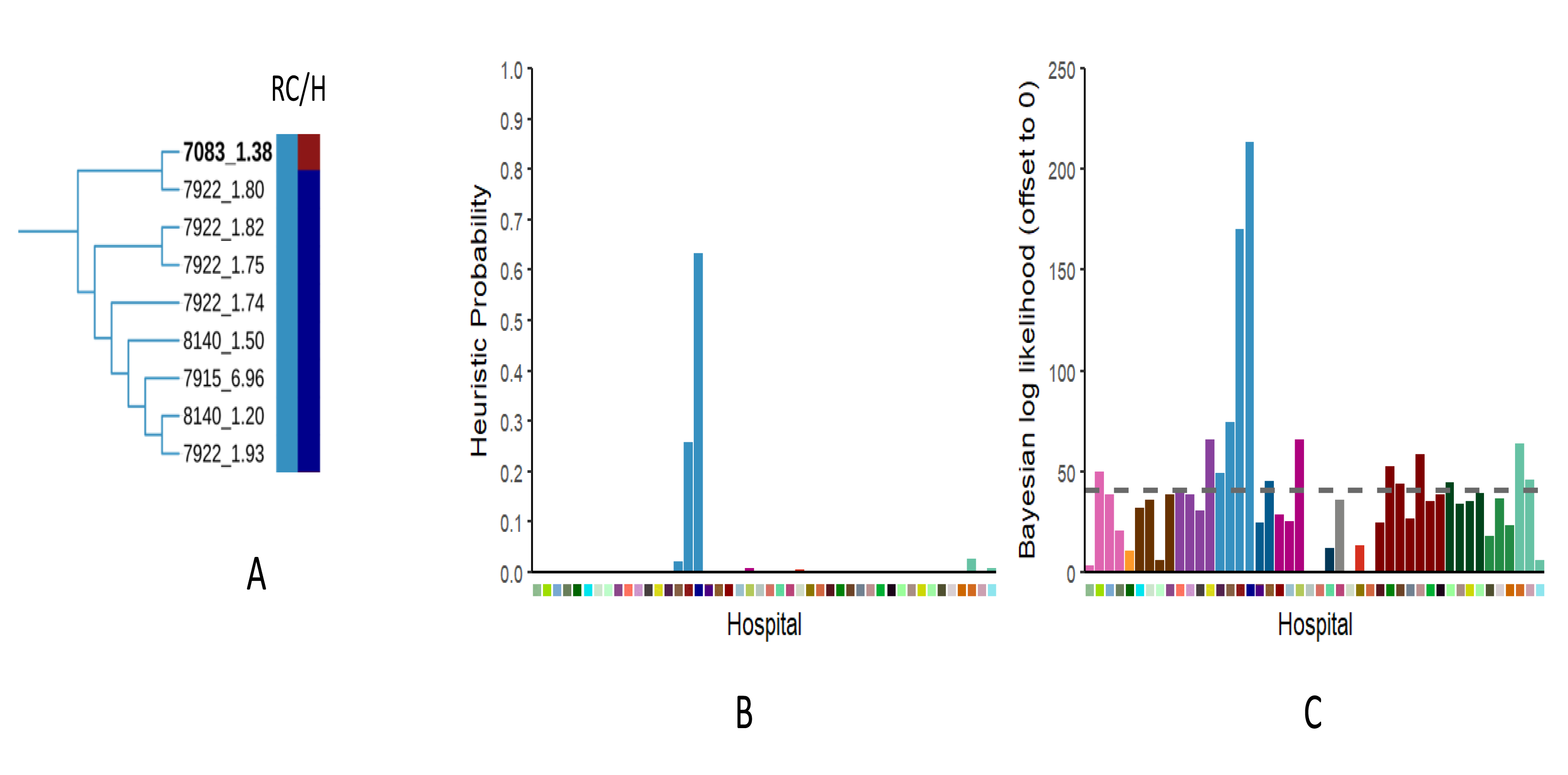
The query isolate id (in bold) is 7083_1.38 was isolated from hospital East2G. However, inspection of the Tree (A) confirms that it corresponds to a clade of isolates from East2D, indicating an inter-hospital transmission event between hospitals belonging to the same referral cluster. Both the heuristic (B) and Bayesian (C) methods predicted East2D as the source of the query isolate, which is consistent with inter-hospital transmission. RC = Referral cluster. H = Hospital. For a key to the colours see Figure 1.

**Figure S6.**
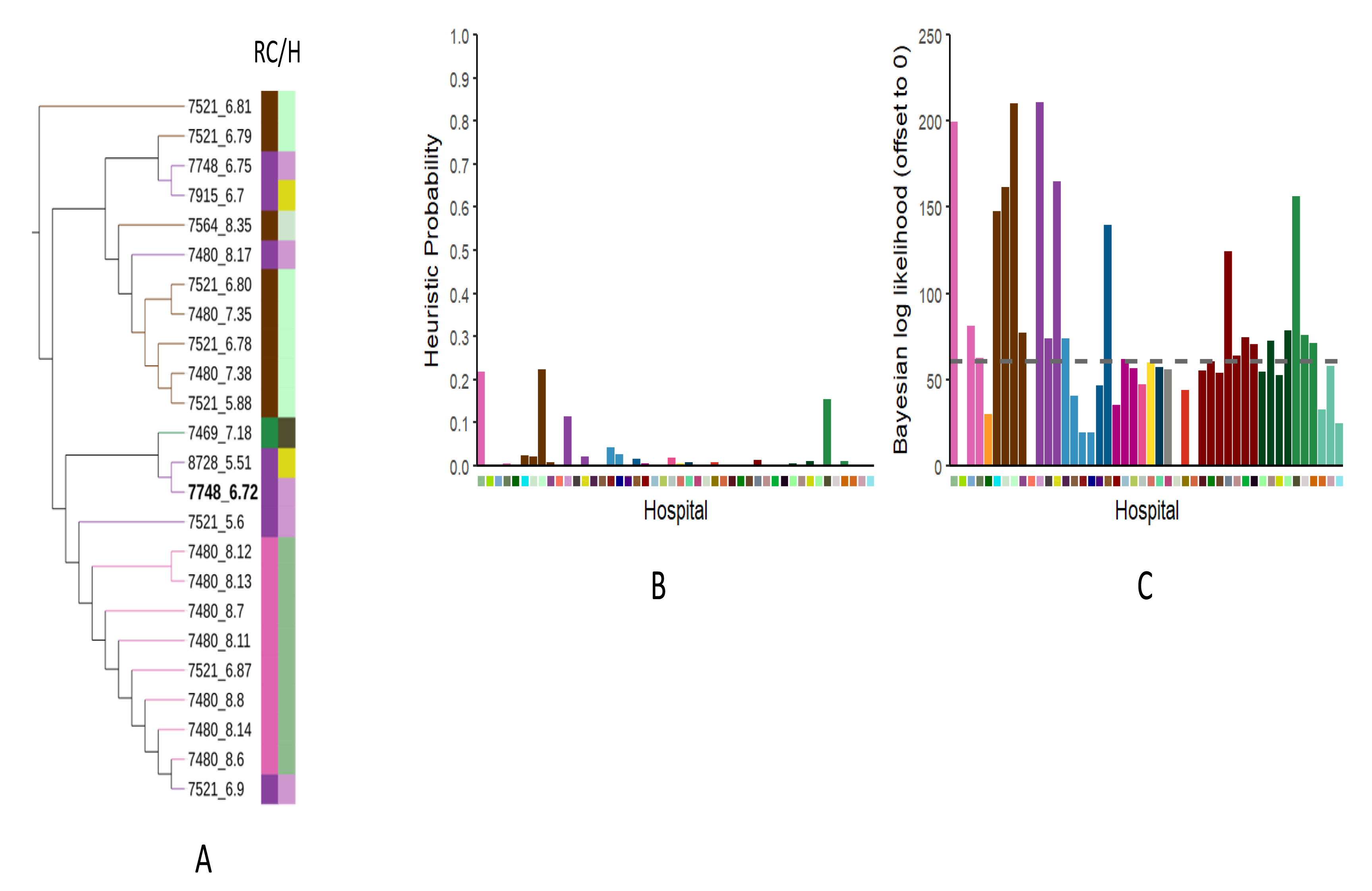
The query isolate id (in bold) is 7748_6.72 was isolated from hospital East1A. However, inspection of the Tree (A) does not provide a strong indication of the source of this isolate. The heuristic (B) predicts the origin as SouthEastC whilst the Bayesian (C) method predicted East1A, which is the actual origin of the isolate. This hospital was favoured in the Bayesian method because of only one other isolate from East1A was present in the database. RC = Referral cluster. H = Hospital. For a key to the colours see Figure 1.

**Figure S7.**
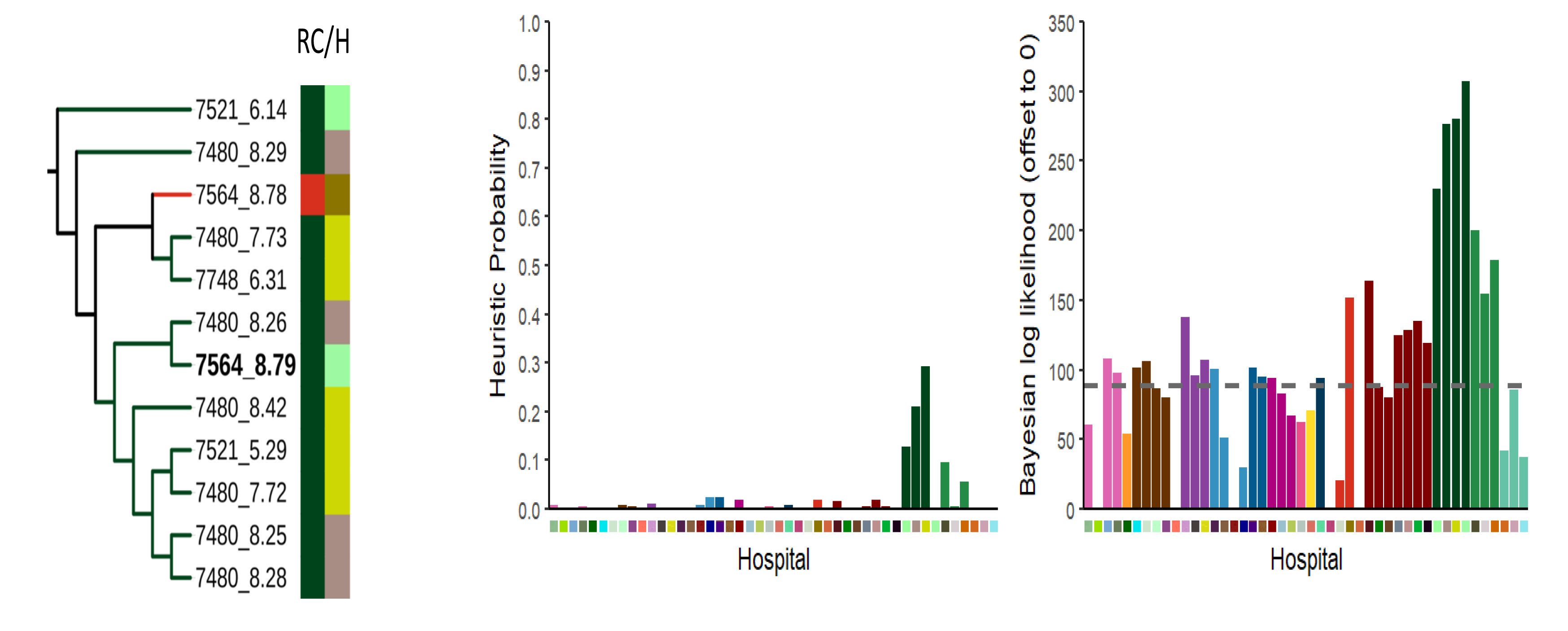
7564_8.79 was isolated from hospital NorthernIrelandB. As this was the earliest isolate recovered from this hospital It was not represented in the database, the heuristic was unable to predict it. However, the Bayseian hospital predicted this hospital due to the prior. RC = Referral cluster. H = Hospital. For a key to the colours see Figure 1.

**Figure S8.**
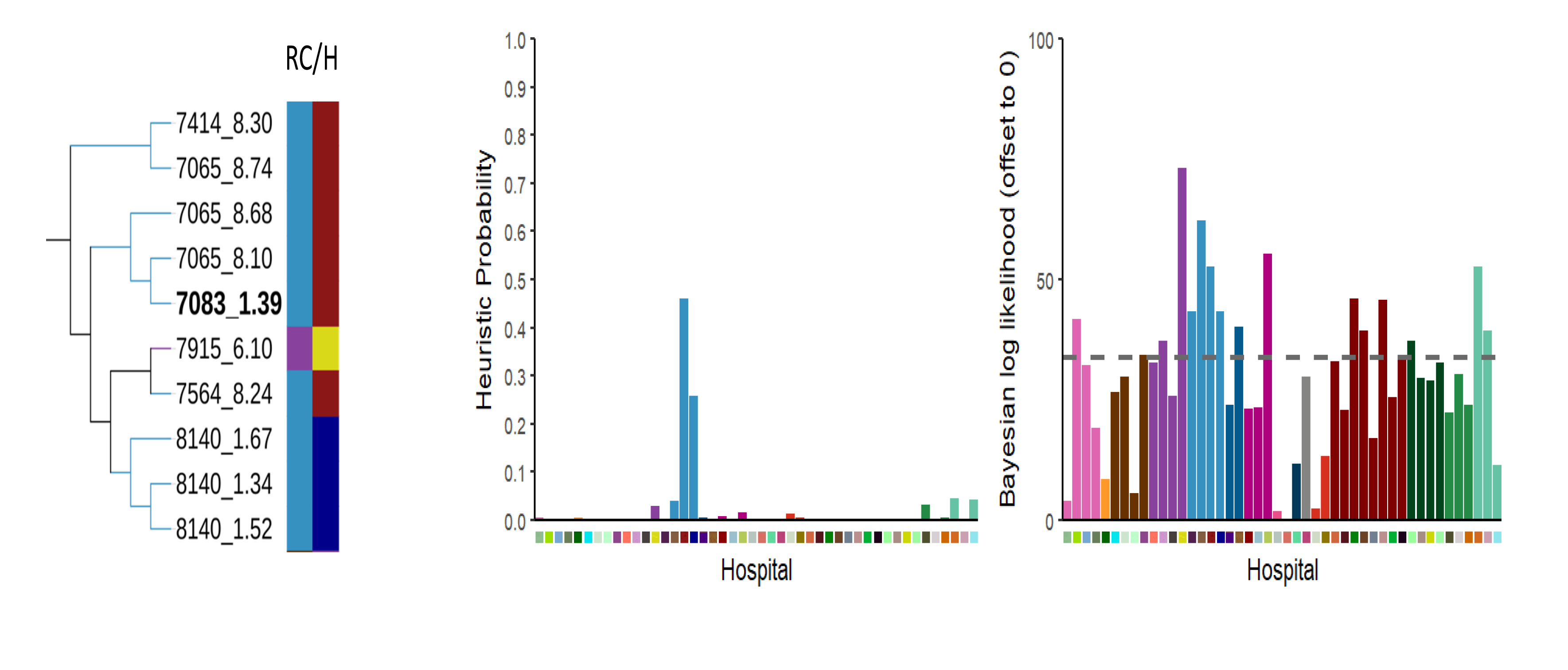
The query isolate id (in bold) is 7083_1.39 was isolated from hospital East2G and the tree (A) shows that it is Related to other isolates from this hospital, pointing to intra-hospital transmission. The heuristic (B) also predicts the origin as East2G. However, the the Bayesian (C) method predicted East1F on the basis that the only isolate corresponding to this hospital in the reference database was reasonably closely related to the query, an that multiple genotypes were present within the well-sample hospital East2G. RC = Referral cluster. H = Hospital. For a key to the colours see Figure 1.

**Figure S9.**
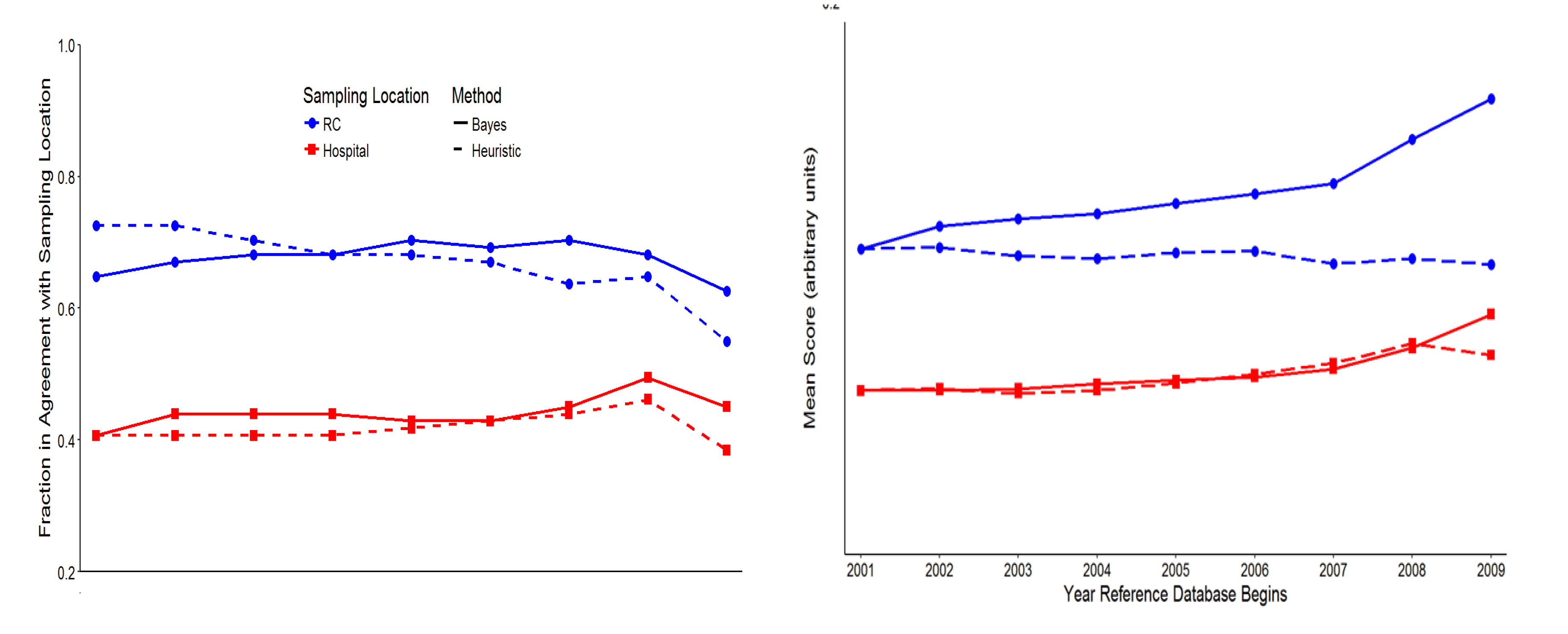
Examining the impact of the temporal breadth of the database. At the hospital level, the percentage of predictions consistent with the actual origin increases slightly at the hospital level as older isolates are removed, but then drops after 2008 presumably due to a paucity of data in the reference database. At the RC level the removal of older isolates has no obvious impact on the Bayesian method, but decreases the consistency of predictions with the actual origin using the heuristic method. Regarding the mean score of the predicted origin, there is a slight increase at the hospital level as older isolates are removed for both methods, but this is more marked when only the later isolates remain for the Bayesian method. There is much more marked increase in the strength of signal at the RC level for the Bayesian method, whilst there is no clear effect using the heuristic method at this level.

**Figure S10.**
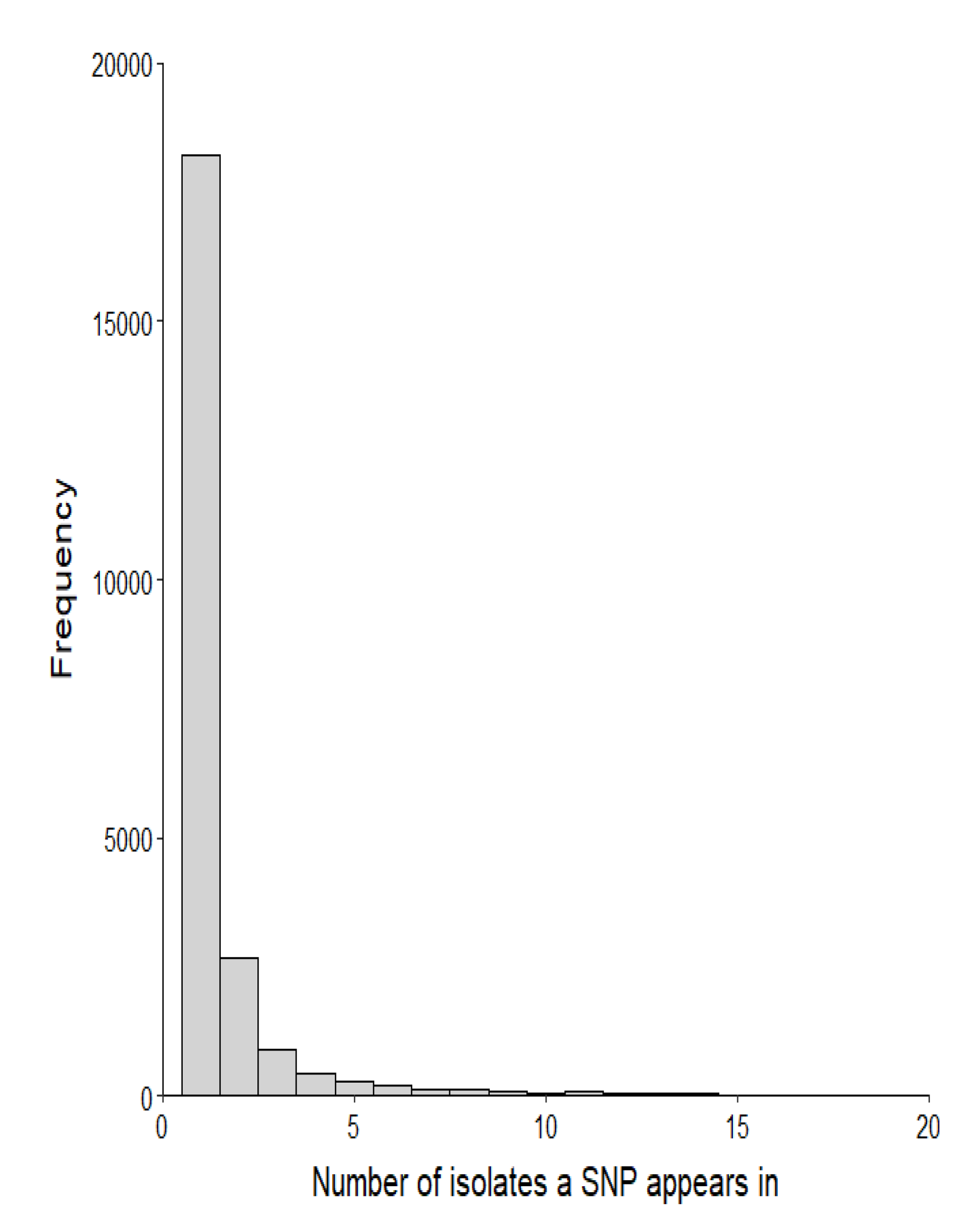
The frequency distribution of SNPs. Data for the very few SNPs found in >20 are not shown

**Figure S11.**
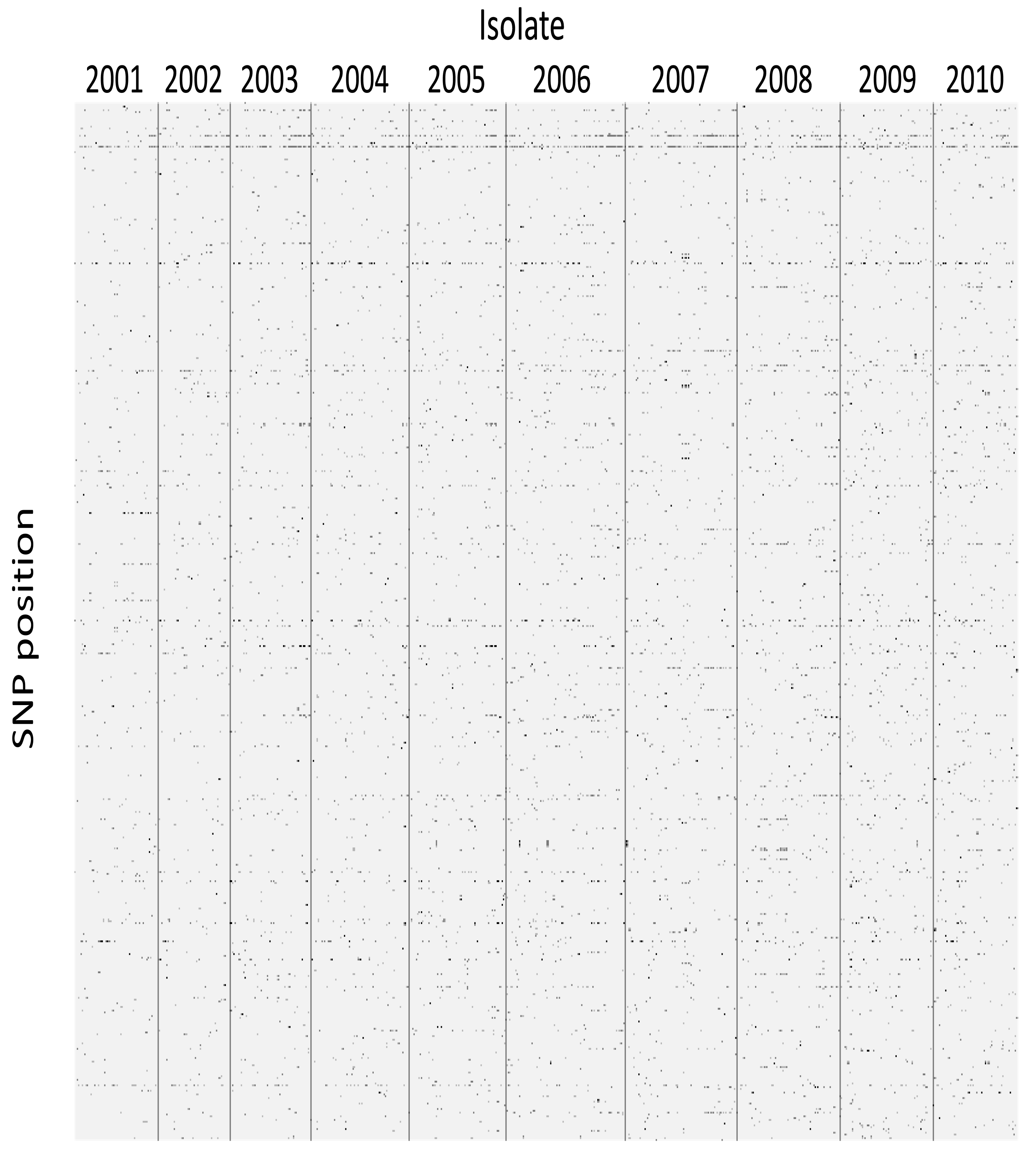
Occurrence of non-singleton SNPs (5469 positions/rows) across the 1022 isolates in the 2001-2010 database. At each position the minority nucleotides are black, and majority nucleotides are grey.

**Figure S12.**
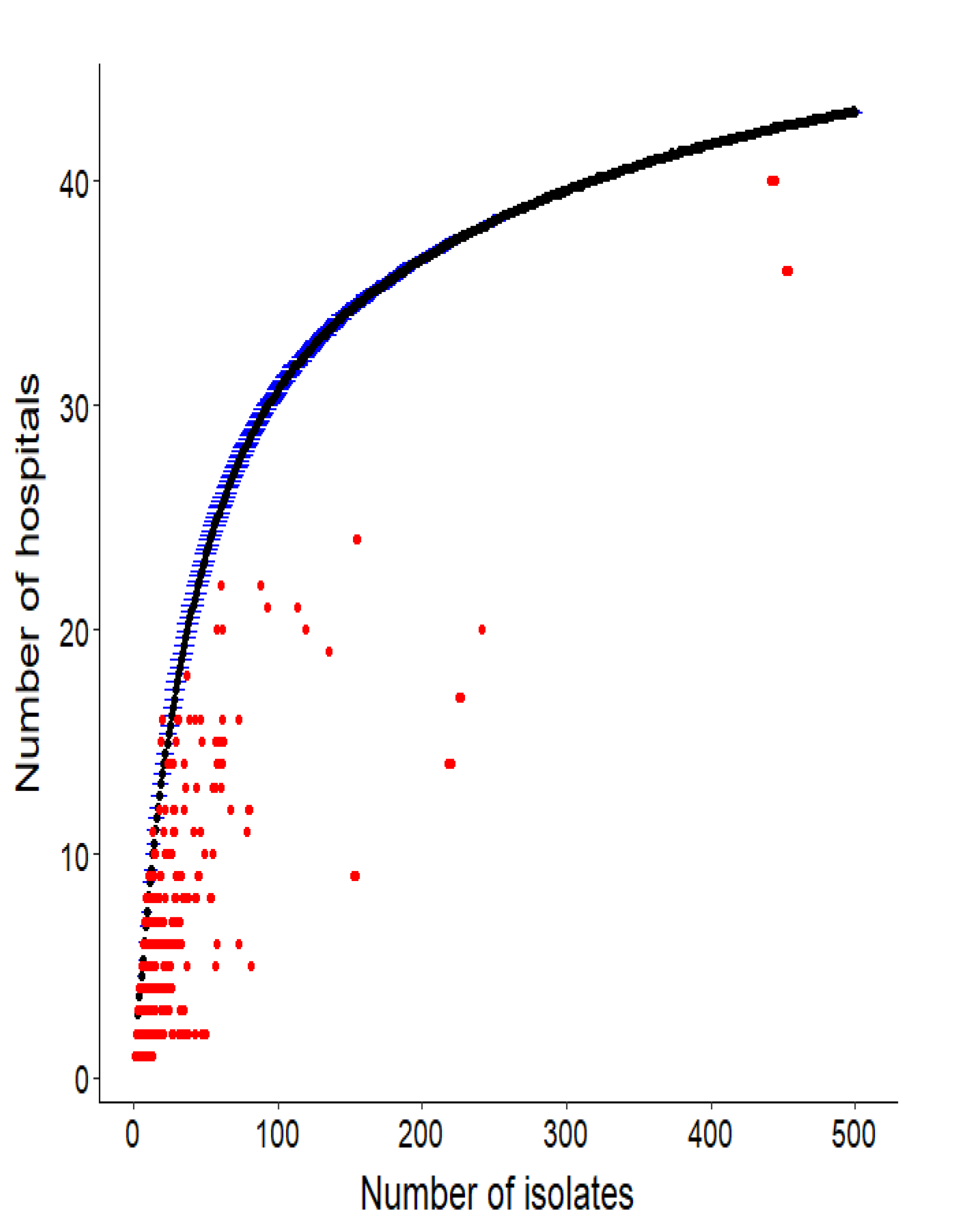
Each point represents a single SNP, and the figure shows, for each SNP, the number of isolates in which that SNP is observed against the number of hospitals. The red points are the observed data, the blue points are expected if the isolates were randomly distributed between hospitals (over 1000 simulations). This shows that SNPs are present in far fewer hospitals than expected, given their frequency in the database.

**Figure S13.**
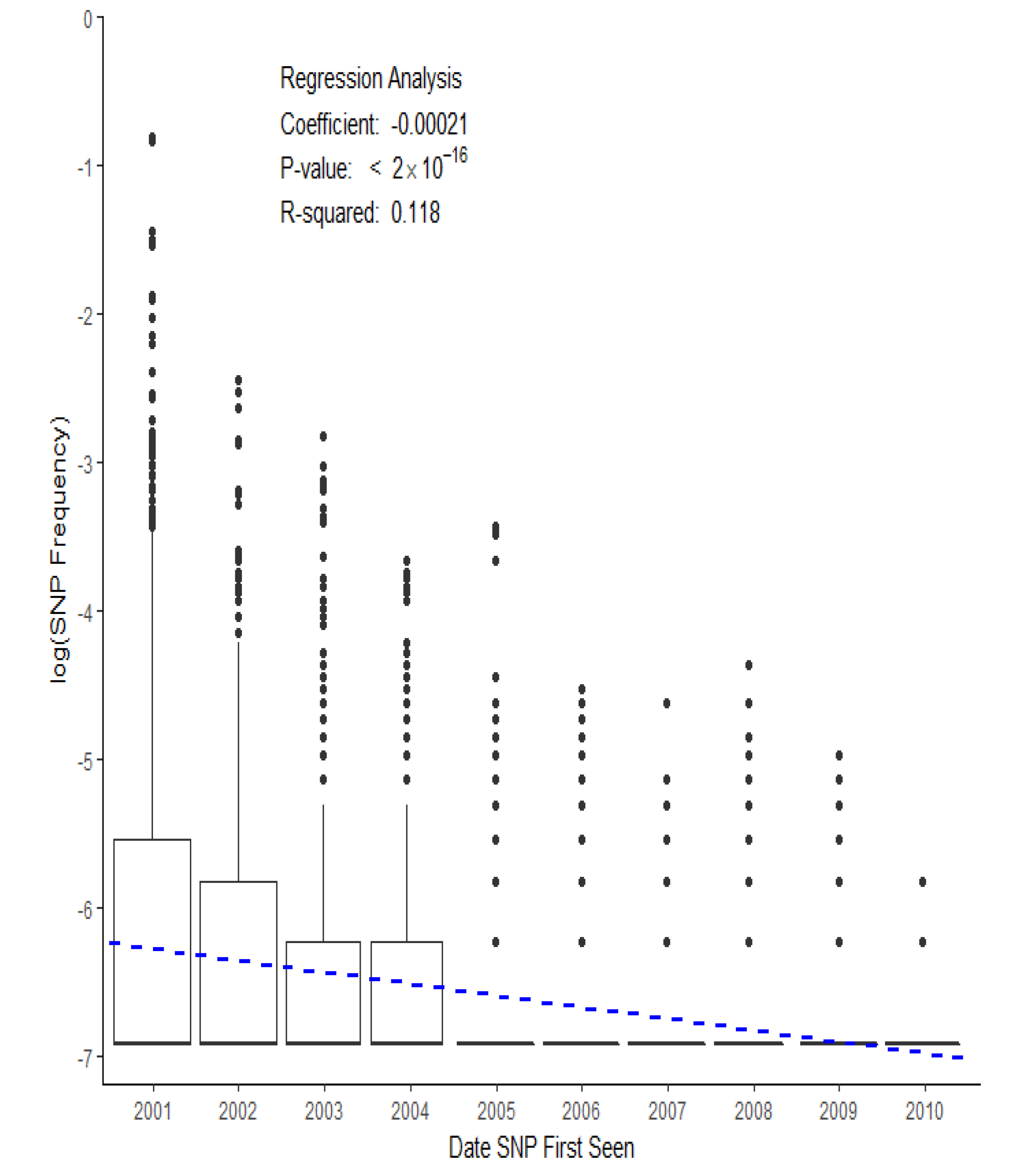
Plot showing relationship between the date at which SNPs are first observed, and their frequency in the database. More recently emerged SNPs are present at a lower frequency and in fewer hospitals, thus they have the greatest discriminatory signal with respect to hospital source.

